# Diverse, abundant and novel viruses infecting “unculturable” but abundant marine bacteria

**DOI:** 10.1101/699256

**Authors:** Zefeng Zhang, Feng Chen, Xiao Chu, Hao Zhang, Haiwei Luo, Fang Qin, Zhiqiang Zhai, Mingyu Yang, Jing Sun, Yanlin Zhao

## Abstract

Many major marine bacterial lineages such as SAR11, *Prochlorococcus*, SAR116, and several *Roseobacter* lineages have members that are abundant, relatively slow-growing, and genome-streamlined. The isolation of phages that infect SAR11 and SAR116 have demonstrated the dominance of these phages in the marine virosphere. However, no phages have been isolated from bacteria in the *Roseobacter* RCA lineage, another abundant group of bacteria in the ocean. In this study, seven RCA phages that infect three different RCA strains were isolated and characterized. All seven RCA phages belong to the *Podoviridae* family and have genome sizes ranging from 39.6 to 58.1 kb. Interestingly, three RCA phages (CRP-1, CRP-2 and CRP-3) show a similar genomic content and architecture with SAR116 phage HMO-2011, which represents one of the most abundant known viral groups in the ocean. The high degree of homology between CRP-1, CRP-2, CRP-3 and HMO-2011 resulted in contribution of the RCA phages to the dominance of HMO-2011-type phage in the ocean. CRP-4 and CRP-5 are similar to the *Siovirus* roseophages in terms of gene content and organization. The remaining two RCA phages, CRP-6 and CRP-7, show limited genomic similarity with known phages and appear to form two new phage genera. Metagenomic fragment recruitment analyses reveal that these RCA phage groups are much more abundant in the ocean compared to most existing marine roseophage groups. The characterization of these RCA phages has greatly expanded our understanding of the genomic diversity and evolution of marine roseophages. Metagenomic fragment recruitment analyses suggest the critical need for isolating phages from the abundant but “unculturable” bacteria in the marine ecosystem.

## Background

Viruses are abundant and infectious to microorganisms in the sea, outnumbering bacteria by an order of magnitude [1, 2]. The majority of marine viruses are bacteriophages whose hosts (bacteria) are the most abundant living organisms in nature [1, 3]. Bacteriophages impact the structure and function of microbial community, and thus have a major influence on the ocean biogeochemical cycles through diverse phage-host interactions [1–3]. Although many marine phages have been isolated, most hosts used for phage isolation are readily-cultivated and fast-growing in rich culture media. For decades, microbial ecologists have been challenged by the fact that a large majority of bacteria in seawater are difficult to be cultivated in the laboratory [4]. Therefore, our understanding on marine virology is greatly hindered by the lack of isolated phages from a broader bacterial groups in the ocean. From the perspective of viromic studies, a major challenge has always been the low percentile of identifiable viral sequences in the viromic databases or viral “dark matter” [5–7]. The large portion of unknown sequences in marine viromes is believed to be due to the lack of known viruses isolated from diverse microbial groups [6, 8]. Despite the gap between viral isolation and viromics, the isolation of some important marine phages, such as SAR116 phage and SAR11 phages (pelagiphages) has greatly facilitated the interpretation of marine viromic datasets [9, 10]. SAR11 and SAR116 are two most abundant and widespread bacterioplankton groups in the ocean [11–13], but the isolates of SAR11 and SAR116 display slow-growth rate and streamlined genomes compared to other cultured representatives from diverse marine bacterioplankton groups [10, 14–16]. The isolation of pelagiphages and SAR116 phage demonstrated that phages infecting abundant but relatively slow-growing marine bacteria make up a significant portion of marine viruses in the ocean [9, 10]. In addition, phage isolation not only provides genomic information, but also morphological and infectious data.

Members of the *Roseobacter* lineage in *Alphaproteobacteria* are abundant, widespread, diverse, and play important biogeochemical roles in the marine environment [17, 18]. Although many roseobacters have been isolated using solid culture media and are readily cultivated in the laboratory, natural *Roseobacter* populations differ systematically from their cultured representatives [19]. Four dominant *Roseobacter* clusters, including *Roseobacter* clade affiliated (RCA, also called the NAC11-3 or DC5-80-3), CHAB-I-5, SAG-O19 and NAC11-7, remain largely uncultivated and poorly studied [20]. These four *Roseobacter* lineages together account for up to >60% of the global pelagic roseobacters [20]. Among these four *Roseobacter* lineages, RCA is the largest cluster of the *Roseobacter* clade that can exceed the SAR11 clade in some oceanic regions [21–23]. RCA is among the most abundant marine bacterioplankton groups, comprising up to 30% of the total bacterioplankton in the temperate and polar ocean regions, and up to 35% of total bacterioplankton with the highest abundance in the Southern Ocean [20–23]. In general, RCA members are difficult to be cultivated using traditional methods. At present, five RCA members have been isolated using the dilution-to-extinction method [21, 24, 25], and only *Planktomarina temperata* RCA23 has been fully sequenced and physiologically characterized [21, 25, 26]. The genome of RCA23 is streamlined, an indication of oligotrophic adaptation [26].

Currently, more than 30 phages that infect representatives of several major *Roseobacter* clusters have been reported [27]. All of these roseophages were isolated from the *Roseobacter* strains that can grow in rich culture media and have fast growth rates, such as *Roseobacter* SIO67, *Ruegeria pomerioyi* DSS-3, *Sulfitobacter* spp. and *Dinoroseobacter shibae* DFL 12 [27–32]. Metagenomic fragment recruitment analyses suggested that homologs of some roseophages were widespread in the ocean, while most of the isolated roseophages were not reported to make up a significant portion of viromic sequences in the ocean. The low abundance of these roseophages is likely associated with low concentration of host cells in the natural environment. The *Roseobacter* lineage contains diverse members, some of which are versatile to environmental changes and can be readily cultured, while others are abundant and often restricted to low nutrient environments, such as the RCA group. Little is known regarding phages that infect the abundant but slow-growing roseobacters [27]. There are multiple challenges associated with isolating phages that infect RCA bacteria. RCA bacteria only grow in diluted media and do not reach high cell densities. Furthermore, they do not grow well in solid media, eliminating the possibility of using plaque assay for phage isolation.

In this study, three different RCA bacteria closely related to *Planktomarina temperata* RCA23 were isolated from the coastal water of Fujian, China. We intended to isolate phages that infect these newly isolated RCA strains. A total of seven RCA phages were isolated and purified using the dilution-to-extinction method. These phages were characterized in terms of their morphology, cross-infectivity, genome sequences, and viromic fragment recruitment. Our results show that RCA phages are abundant and diverse in the ocean, and contain many genome types that were not previously recognized.

## Results and discussion

### Host strains

Three *Roseobacter* strains, FZCC0023, FZCC0040, and FZCC0042, were isolated in 2017 from the coastal water of Pingtang Island, Fujian, China using the dilution-to-extinction method [16]. Phylogenetic analysis based on the 16S rRNA gene sequences suggests that FZCC0023, FZCC0040, and FZCC0042 all belong to the RCA cluster (Additional file 1: Figure S1a). In terms of the nearly complete 16S rRNA gene sequence, FZCC0040 and FZCC0042 are 100% identical to RCA23, while FZCC0023 has 2 nucleotide mismatches with RCA23. However, these RCA strains can be distinguished from RCA23 and each other based on their 16S-23S rDNA intergenic spacer (ITS) sequences, suggesting that they are closely related but distinct RCA members (Additional file 1: Figure S1b).

### Isolation and general features of RCA phages

Seven phages (CRP-1, CRP-2, CRP-3, CRP-4, CRP-5, CRP-6 and CRP-7) that infect the three above-mentioned RCA strains (hereafter referred to as RCA phages) were isolated from different locations ranging from temperate (Bohai Sea, China and Osaka Bay, Japan) to subtropical (Pingtang coast, China) regions (Table 1). All seven RCA phages have short tails and icosahedral capsids (Fig. 1a); thus, they all belong to the *Podoviridae* family in the order *Caudovirales*. Except for CRP-7, the RCA phages did not cross infect other RCA strains beyond the original host (Table 1). CRP-7, which was originally isolated from FZCC0042, was also able to infect FZCC0040. Considering the high 16S rRNA gene sequence identity of the tested RCA strains, this result suggests that these RCA phages appear have a very narrow host range. The complete genomes of these seven RCA phages were sequenced and assembled, and their genome sizes vary from 39 to 55 kb. The G+C content of these phages ranges from 40.3 to 49.7%, lower than that of their hosts (50 to 53%). The RCA phages appear to have a relatively lower G+C content compared to other sequenced roseophages (43 to 64%) [27]. Except for CRP-7, the RCA phages do not contain tRNA sequences. The predicted ORFs are provided in Dataset 1 (Additional file 3). Comparative genomics analysis categorized these 7 RCA phages into four distinct phage genera within the *Podoviridae* family (Fig. 1b and Fig. 2). Taken together, these results suggest that RCA roseobacters are subjected to phage infection by diverse podoviruses (non N4-like podoviruses), a scenario different from the known roseophages that infect readily cultivated marine roseobacters [27]. Among the 32 known roseophages, siphoviruses and N4-like podoviruses dominate the current isolates [27].

**Fig. 1.**
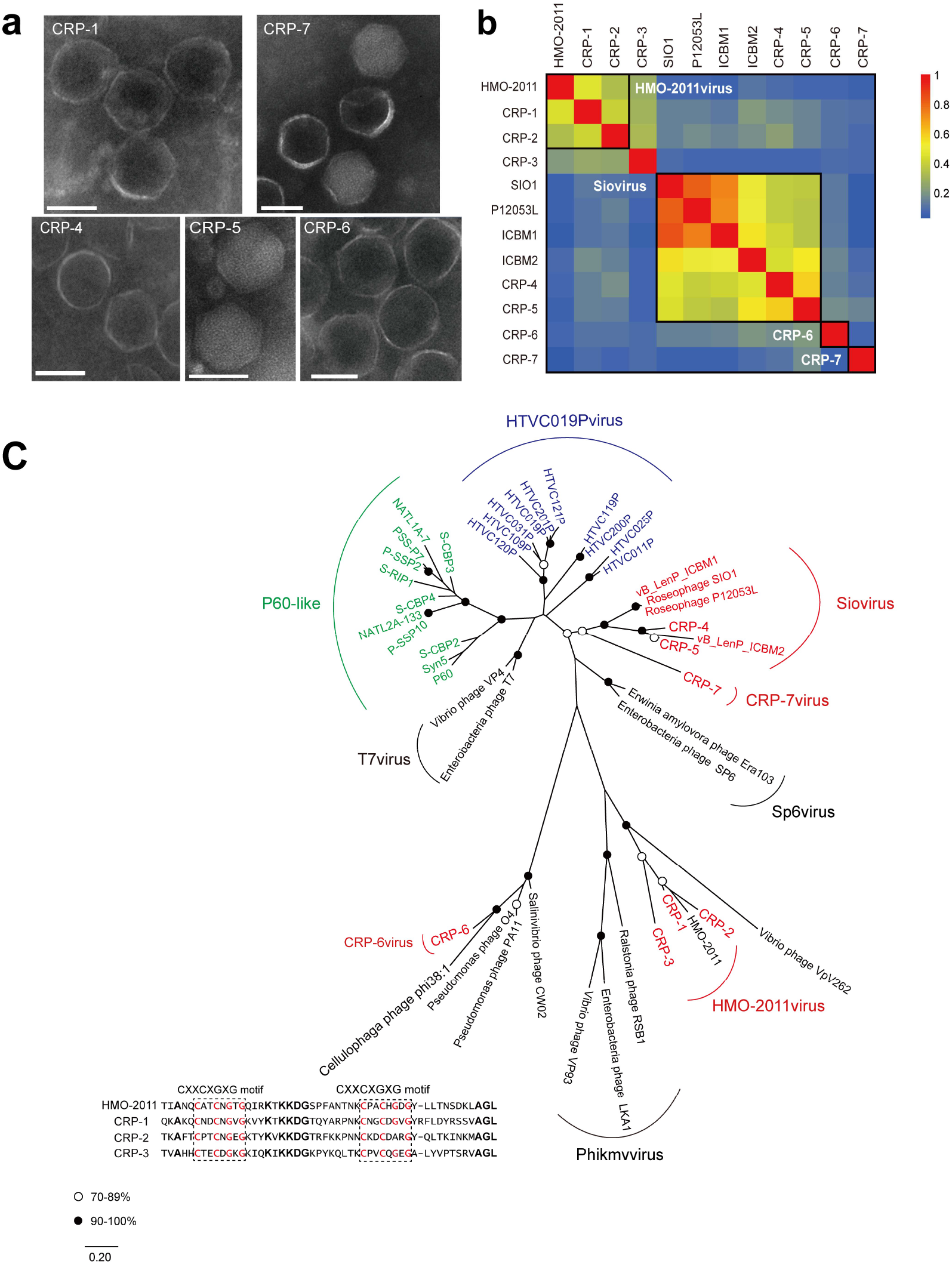
**a** Transmission electron microscopy images of selected representative RCA phages from each genus (Scale bars: 50 nm.). **b** Heatmap presentation of shared genes of newly isolated RCA phages and five related marine phages (HMO-2011, SIO1, P12053L, ICBM1 and ICBM2). The genome similarity between the phages refers to the heatmap bar on the right. Phages in the same genus are boxed. **c** Unrooted maximum-likelihood phylogenetic tree of phage DNA polymerases constructed with conserved polymerase domains. Black and white circles indicate nodes with 90-100% and 70-89% bootstrap support, respectively. Roseophages, cyanophages and pelagiphages are shown in red, green and blue, respectively. Scale bar represents amino acid substitutions per site. The alignment of the partial sequence of DnaJ central domain from HMO-2011 and three RCA phages is presented. Two CXXCXGCG motifs are boxed. Conserved residues in the two CXXCXGXG motifs are in red. Other conserved residues are in bold.

**Fig. 2.**
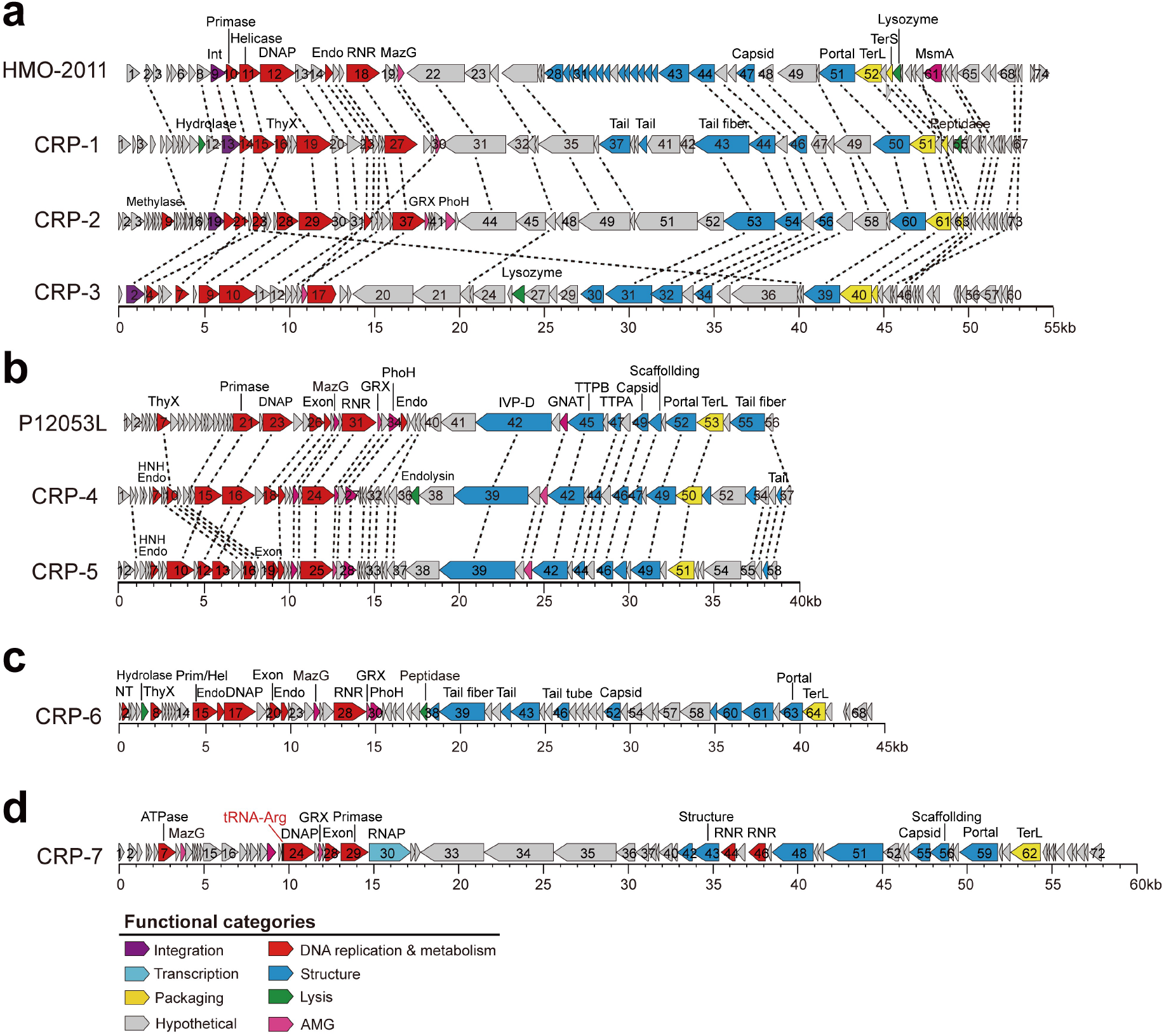
Genomic organization and comparison of RCA phages. **a** *HMO-2011virus* phages and CRP-3. **b** *Siovirus* roseophages. **c** CRP-6. **d** CRP-7. ORFs are depicted by leftward or rightward oriented arrows according to their transcription direction. ORFs are color-coded according to putative biological function. Homologous ORFs are connected by dash lines. ORFs homologous to HMO-2011 are connected by red dash lines. tRNAs are shown in red. Abbreviation: int, integrase; RNA pol, RNA polymerase; SSB, single-stranded DNA binding protein; endo, endonuclease; DNAP, DNA polymerase; exon, exonuclease; TerS, terminase, small subunit; TerL, terminase, large subunit. ThyX, thymidylate synthase; MazG, MazG nucleotide pyrophosphohydrolase domain protein; RNR, adenosylcobalamin-dependent ribonucleoside-triphosphate reductase; GNAT, GCN5-Related N-Acetyltransferases Acetyltransferase family; IVP-D, internal virion proteins D; TTPA, Tail tubular protein A; TTPB, Tail tubular protein B; GRX, glutaredoxin; NT, Nucleotidyltransferase;

**Table 1.**
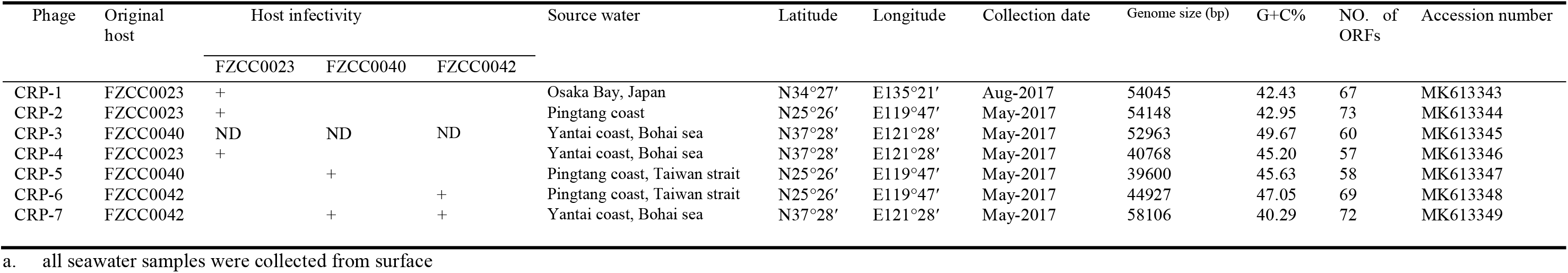
General features of seven RCA phages analyzed in this study

| Phage | Original host | Host infectivity |  |  | Source water | Latitude | Longitude | Collection date | Genome size (bp) | G+C% | NO. of ORFs | Accession number |
| --- | --- | --- | --- | --- | --- | --- | --- | --- | --- | --- | --- | --- |
|  |  | FZCC0023 | FZCC0040 | FZCC0042 |  |  |  |  |  |  |  |  |
| CRP-1 | FZCC0023 | + |  |  | Osaka Bay, Japan | N34°27' | E135°21' | Aug-2017 | 54045 | 42.43 | 67 | MK613343 |
| CRP-2 | FZCC0023 | + |  |  | Pingtang coast | N25°26' | E119°47' | May-2017 | 54148 | 42.95 | 73 | MK613344 |
| CRP-3 | FZCC0040 | ND | ND | ND | Yantai coast, Bohai sea | N37°28' | E121°28' | May-2017 | 52963 | 49.67 | 60 | MK613345 |
| CRP-4 | FZCC0023 | + |  |  | Yantai coast, Bohai sea | N37°28' | E121°28' | May-2017 | 40768 | 45.20 | 57 | MK613346 |
| CRP-5 | FZCC0040 |  | + |  | Pingtang coast, Taiwan strait | N25°26' | E119°47' | May-2017 | 39600 | 45.63 | 58 | MK613347 |
| CRP-6 | FZCC0042 |  |  | + | Pingtang coast, Taiwan strait | N25°26' | E119°47' | May-2017 | 44927 | 47.05 | 69 | MK613348 |
| CRP-7 | FZCC0042 |  | + | + | Yantai coast, Bohai sea | N37°28' | E121°28' | May-2017 | 58106 | 40.29 | 72 | MK613349 |

a. all seawater samples were collected from surface

### Close kinship between RCA phages and SAR116 phage HMO-2011

Three RCA phages (CRP-1, CRP-2 and CRP-3) share a similar genomic content and arrangement with a recently sequenced phage, HMO-2011, which infects a SAR116 bacterium “*Candidatus* Puniceispirillum marinum” strain IMCC1322 (Fig. 2a). CRP-1, CRP-2 and CRP-3 have 30, 25 and 16 ORFs homologous to genes in the HMO-2011 genome, respectively. About half of the shared genes in CRP-1 and CRP-2 are >50% identical in amino acid sequences to their counterparts in HMO-2011, while most shared genes in CRP3 are <50% identical in amino acid sequences to their HMO-2011 counterparts. The shared genes are primarily involved in the essential functions for the phage life cycle including DNA metabolism and replication, phage structure and DNA packaging, and they are arranged in a conserved order (Fig. 2a). There is no significant genomic feature that distinguishes these three RCA phages from SAR116 phage. Only three genes were observed to be exclusive to these three RCA phages, including a thymidylate synthase gene, a tail fiber gene and a gene encoding an unknown function protein (ORF47 in CRP-3). Phage tail fibers are responsible for host specificity. The variation in tail fiber genes between the RCA phages and the SAR116 phage suggests an adaptation of phages to different host systems. Based on the criteria of >40% and >20% of shared genes for genus and subfamily discrimination [33], respectively, CRP-1, CRP-2 and HMO-2011 can be grouped into a genus that was designated as the putative *HMO-2011virus* genus. CRP-3 is more distantly related and can be classified with CRP-1, CRP-2 and HMO-2011 at the subfamily-level (Fig. 1b). Although HMO-2011-type phages is among the most abundant known viral groups in the ocean [10], HMO-2011 has no counterparts among currently known phages. The high genomic homology among CRP-1, CRP-2, CRP-3 and HMO-2011 suggests a close kinship between the RCA phages and the SAR116 phage.

The close relationship between the three RCA phages and HMO-2011 is also evident based on the DNA polymerase gene, a gene that is particularly conserved among marine podoviruses [34]. The amino acid sequences of the DNA polymerases of these three RCA phages are 40 to 67% identical to that of HMO-2011. The DNA polymerase gene phylogeny shows that CRP-1, CRP-2 and HMO-2011 cluster into one group, while CRP-3 forms its own branch adjacent to this group (Fig. 1c). The HMO-2011 DNA polymerase possesses an unusual domain architecture, with a partial DnaJ domain located between the exonuclease domain and the DNA polymerase domain [10]. The DNA polymerases of these three RCA phages also exhibit this unusual domain structure, and contain two CXXCXGXG motifs in the partial DnaJ domain (See the box in Fig. 1c). CRP-1, CRP-2 and CRP-3 all encode a tyrosine integrase upstream of the DNA replication and metabolism module, which shares 35 to 63% amino acid identity with the HMO-2011 integrase. Integrase genes typically occur in the genomes of temperate phages and are responsible for site-specific integration. The presence of an integrase suggests that these phages possibly undergo a lysogenic life cycle. We demonstrated that CRP-3 can integrate into a tRNA-Met (CAT) site in the FZCC0040 genome (Additional file 1: Figure S2a), suggesting that CRP-3 reproduction occurs via lytic and lysogenic cycles. The core sequence overlaps the 3’ end of the host tRNA-Met gene (Additional file 1: Figure S2b). However, the integration sites of CRP-1 and CRP-2 in FZCC0023 have not yet been identified; thus, it is still unknown whether CRP-1 and CRP-2 also have a lysogenic life cycle.

These results suggest that HMO-2011-type phage (at the subfamily-level) can infect a broad range of bacterial hosts. It is interesting that within the *HMO-2011virus* genus, members infecting SAR116 and RCA roseobacters are strikingly similar. The high sequence homology between the RCA phages and SAR116 phages raises a concern on potential overlaps among these phages on viromic fragment recruitment (see the later section Viromic fragment recruitment analyses of RCA phages)

It is general observed that phages infecting the closely related hosts appeared to be more closely related and a recent phage phylogeny analysis suggested that phage genera usually infect bacteria within the same family [35]. By contrast, in our study, closely related *HMO-2011virus* phages were observed to infect both SAR116 and RCA bacteria, which belong to two distinct orders. SAR116 and RCA both possess high population densities [12, 13, 21, 22, 36], are therefore among the most common phage hosts in the ocean. In addition, SAR116 and RCA display a similar distribution pattern in the global ocean, are both predominate in temperate and polar ocean [13, 22, 36]. Considering the high population densities of SAR116 and RCA, our results imply that common ancestors of these phages were more likely to collide with these abundant bacteria by chance and evolved to gain the ability to attach and take control of host machinery. Thus, the genome sequences of these RCA phages provide important clues for understanding the evolution and taxonomy of this important phage group.

### Two RCA phages are closely related to *Siovirus* roseophages

CRP-4 and CRP-5 share similar genome content and architecture with the roseophages SIO1, P12053L, ICBM1 and ICBM2, which infect *Roseobacter* SIO, *Celeribacter* sp. strain IMCC12053 and *Lentibacter* sp. SH36, respectively [28, 37, 38] (Fig. 2b). Although these roseophages are related to the phages of the *Autographivirinae* subfamily from an evolutionary perspective, they were previously classified as an unassigned *Podoviridae* group because they all lack a phage-encoded RNA polymerase [33]. Recently a new genus *Siovirus* (within the *Riovirinae* subfamily) was established after the isolation of three marine cobaviruses (cobalamin-dependent) [38]. CRP-4 and CRP-5 also lack a RNA polymerase gene (Fig. 2b), and they should be classified as new members of the *Siovirus* genus based on their genome synteny (Fig. 1b and Fig. 2b). Most of the shared genes within the *Siovirus* genus are located in the DNA replication module, including genes predicted to encode thymidine synthase, a DNA primase, a T7-like DNA polymerase, an endonuclease and a ribonucleotide reductase. Other homologous genes encode proteins involved in the phage virion structure and DNA packaging, such as the coat protein, portal protein, tail proteins, and large subunit of terminase (Fig. 2b). The DNA polymerase gene phylogeny indicates that CRP-4 and CRP-5 are clustered with four other *Siovirus* roseophages, as expected (Fig. 1c). *Siovirus* roseophages are further separated into two well-supported subclusters, with CRP-4 and CRP-5 clustering with ICMB2, SIO1 and P12053L forming a distinct subcluster. (Fig. 1c).

### CRP-6 and CRP-7 represent two novel phage groups

CRP-6 shares limited homology with other known phage isolates, it can therefore be classified into the putative *CRP-6virus* genus in the *Podoviridae* family. Nearly 30% of the ORFs in CRP-6 can be assigned putative biological functions (Fig. 2c). Although CRP-6 appears to be closely related to *Cellulophaga* phage phi38:1 and *Salinivibrio* phage CW02 based on the DNA polymerase phylogeny (Fig. 1c), CRP-6 only shares few genes with phi38:1 and CW02 at the genomic level. Therefore, in this case, the DNA polymerase gene phylogeny does not reflect the genomic evolution of CRP-6. The genome of CRP-6 appears to be highly mosaic, as it shares the DNA replication genes with some podoviruses, but shares its structural genes with other types of phages. For example, the CRP-6 primase/helicase gene is most closely related to those found in *Siovirus* roseophages; the DNA polymerase gene of CRP-6 is mostly related to that of *Cellulophaga* phage phi38:1; the capsid gene of CRP-6 is similar to that of roseosiphovirus RDJL Phi 1; and the tail fiber gene of CRP-6 is similar to that of cyanomyovirus S-CAM7. CRP-6 encodes a terminase large subunit-like protein with homologs found in some bacterial genomes.

CRP-7 has the largest genome size among the seven RCA phage isolates (Table 1). The CRP-7 genome is 58.1 kb in length, consisting of 73 predicted ORFs and a tRNA-Arg (TCT) gene. Similar to CRP-6, CRP-7 also exhibits no significant genomic synteny with other known phage isolates. Therefore, we propose a new genus, the putative *CRP-7virus* in the *Podoviridae* family. Of the 73 predicted ORFs, 35 have homologs in the NCBI-RefSeq database and only 18 have assigned functions (Fig. 2d). Functional predictions for the annotated ORFs were predominantly associated with phage DNA replication and virion morphogenesis. CRP-7 contains a suit of DNA replication genes with divergent similarity to the phages in the *Podoviridae* family. The DNA polymerase of CRP-7 encoded by ORF24 is mostly related to the DNA polymerase in *Siovirus* roseophages. ORF 30 was predicted to encode a DNA-dependent RNA polymerase, having distant relationship to the RNA polymerase in members of the *Autographivirinae* subfamily (<25% amino acid identity). Additionally, several structural genes were predicted from the CRP-7 genome, most of which show weak homology to structural proteins in other *Podoviridae* phages. The gene encoding terminase large subunit in CRP-7 is also very divergent, showing similarity with some bacterial terminases genes and distant relationship to the terminases in some T4-like cyanomyoviruses. The DNA polymerase gene phylogeny reveals that CRP-7 is distantly related to other known marine podoviruses, forming a *CRP-7virus* branch near *Siovirus* roseophages (Fig. 1c).

### Novelty of the RCA phages

We built a gene-content-based network to illustrate the relationship of RCA phages to other phages in the *Podoviridae* family (list in Additional files2: Table S1). The seven RCA phages are grouped into four distinct clusters (Fig. 3). In agreement with the genomic comparative analysis and the DNA polymerase gene phylogeny, CRP-1, CRP-2 are grouped with HMO-2011, belonging to the *HMO-2011virus* genus. CRP-3 is more distantly related to phages in *HMO-2011virus* genus. CRP-4 and CRP-5 cluster with *Siovirus* roseophages, while CRP-6 and CRP-7 are both distantly related to other podoviruses which infect *Roseobacter*. All four RCA phage groups show weak relationships to the well-known phages in the *Autographivirinae* subfamily and some other phages in the *Podoviridae* family. The network analysis further supports that phages infecting RCA can be diverse and novel to our current collection of marine phages (Fig. 3).

**Fig. 3.**
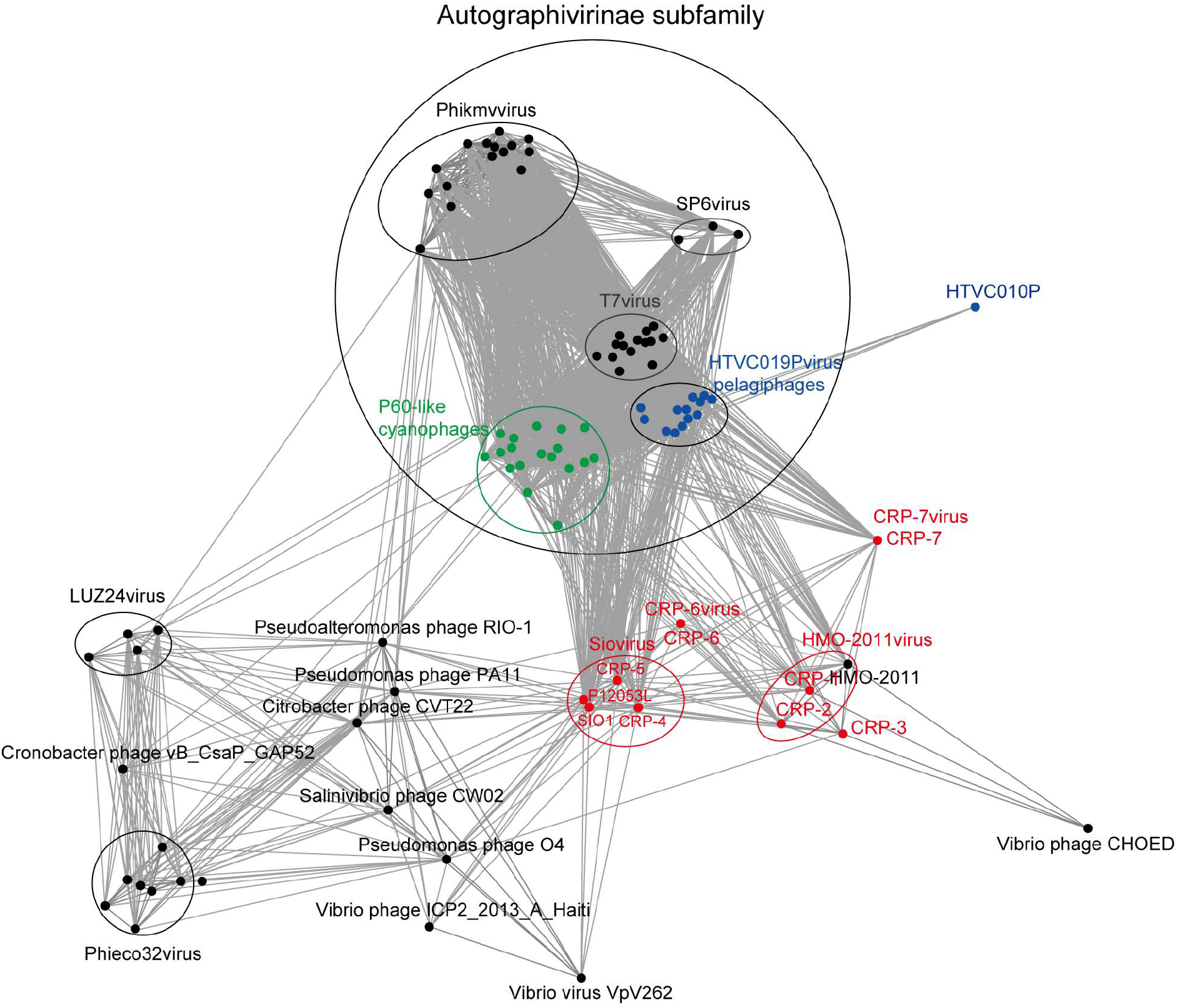
Gene-content-based viral network of RCA phages and representative *podoviridae* phages. Roseophages, pelagiphages and cyanophage are represented as red, blue and green nodes, respectively. The edges represent phage–phage similarities in terms of shared gene ≥0.05. Different viral genera or groups are circled.

Interestingly, all seven RCA phages belong to the *Podoviridae*. It is somewhat surprising that RCA podoviruses show greater genomic diversity compared to the cyanopodoviruses that infect marine *Synechococcus* and *Prochlorococcus* [39]; considering that three RCA host strains we used are closely related. Similarly, high genomic diversity was also reported on marine *Cellulophaga* podophages isolated using closely related hosts [40].

### RCA phage auxiliary metabolic genes and genes involved in other cellular processes

All RCA phage genomes possess several auxiliary metabolic genes (AMGs), which are presumably of bacterial origin. AMGs play roles in the regulation of host metabolism during host infection and therefore benefit phage production [41]. The AMGs identified from the seven RCA phage genomes are summarized in Table S2 (Additional files2). All seven RCA phages harbour a gene encoding an adenosylcobalamin-dependent ribonucleoside-triphosphate reductase (RNR) (PF02867.14) and all but one RCA phage harbour a gene encoding a thymidylate synthase, which is possibly involved in phage nucleotide metabolism. Another AMG involved in nucleotide metabolism is a putative DNA (cytosine-5) methylase gene identified from CRP-2. DNA methylase is involved in cytosine residue methylation commonly occurs in bacterial genomes and has been also found in many phage genomes. This DNA methylase may possibly methylate the phage sequence and protect it against host restriction-modification (R-M) systems [42].

Five RCA phages invistigated in this study also possess a starvation-inducible protein gene (*phoH*), suggesting its importance in the phosphate metabolism of phage-infected roseobacterial cells.

All but one RCA phage encodes a conserved the MazG pyrophosphohydrolase domain (PF03819). The MazG gene was also identified in many marine phage genomes [10, 43–45]. MazG proteins are involved in the regulation of bacterial survival under nutritional stress [46]. MazG proteins can extend the period of cell survival, which is important for phage reproduction [47]. It was hypothesized that phages encoded MazG proteins play an important role in phage propagation by maintaining host metabolism under stress [10, 43, 45].

Glutaredoxins genes were predicted from five RCA phage genomes. Glutaredoxins are redox proteins that have been shown to be involved in multiple cellular processes, such as oxidative stress response, deoxyribonucleotide synthesis, repair of oxidatively damaged proteins, protein folding, sulphur metabolism and other redox-dependent signaling, by catalyzing glutathione-disulfide oxidoreductions [48]. Glutaredoxins also serve as hydrogen donors in the redox reactions of the RNR catalysis [49]. It was suggested that glutaredoxins may contribute to phage propagation by maintaining the cellular redox state of the host and by interacting with the phage-encoded ribonucleotide reductase [49].

Interestingly, CRP-4 and CRP-5 each encode a GCN5-related N-acetyltransferases (GNAT). GNATs catalyze the transfer of an acyl group from acyl coenzyme A (acyl-CoA) to an amino group of a wide range of substrates [50]. GNATs are involved in diverse cellular processes, including carbohydrate and energy metabolism, nucleotide and amino acid metabolism, transcription, translation, cell differentiation, stress regulation and many others [50]. Because GNATs are integral to bacterial metabolism, it is thus suggested that CRP-4 and CRP-5 may regulate primary host metabolism to complete phage propagation through GNAT [50].

CRP-6 contains a gene encoding glycerol-3-phosphate cytidylyltransferase (GCT) (PF01467.25 CL0039), which is involved in the biosynthesis of teichoic acid and is required for bacterial cell wall biogenesis [51]. The function of the GTC-encoding gene in CRP-6 remains unclear.

### Viromic fragment recruitment analyses of RCA phages

Metagenomic fragment recruitment analyses were performed to assess the distribution and relative abundance of RCA phages compared with other important marine phage groups and phage isolates. A total of 174 marine viromic datasets that from the Pacific Ocean Virome (POV), Scripps Pier Virome (SPV), India Ocean Virome (IOV), Malaspina Expedition virome (ME) and Global Oceans Viromes (GOV) were used for the recruitment analyses (Additional files2: Table S3), which cover a wide range of marine habitats. The HMO-2011-type phage group (at the subfamily-level, here including *HMO-2011virus* phages and CRP-3) was the most abundant known phage group in most of the marine viromes (Fig. 4, Additional file 1: Figure S6). These data are consistent with the previous finding that the phage type represented by HMO-2011 is among the most abundant viral groups in marine viromes [10]. We also noticed that the reads assigned to CRP-3 only account for approximately 10% of the total reads assigned to this group (data not shown), suggesting that phages closely related to CRP-3 are not a dominant type. Our results explain the high abundance of the HMO-2011-type group in the ocean, as phages in this group could infect diverse groups of bacterial hosts. Due to the high sequence homology between the RCA phages and HMO-2011, it is obvious that the RCA phages, and probably other undiscovered marine phages also contribute to the abundance and diversity of the HMO-2011-type group.

**Fig. 4.**
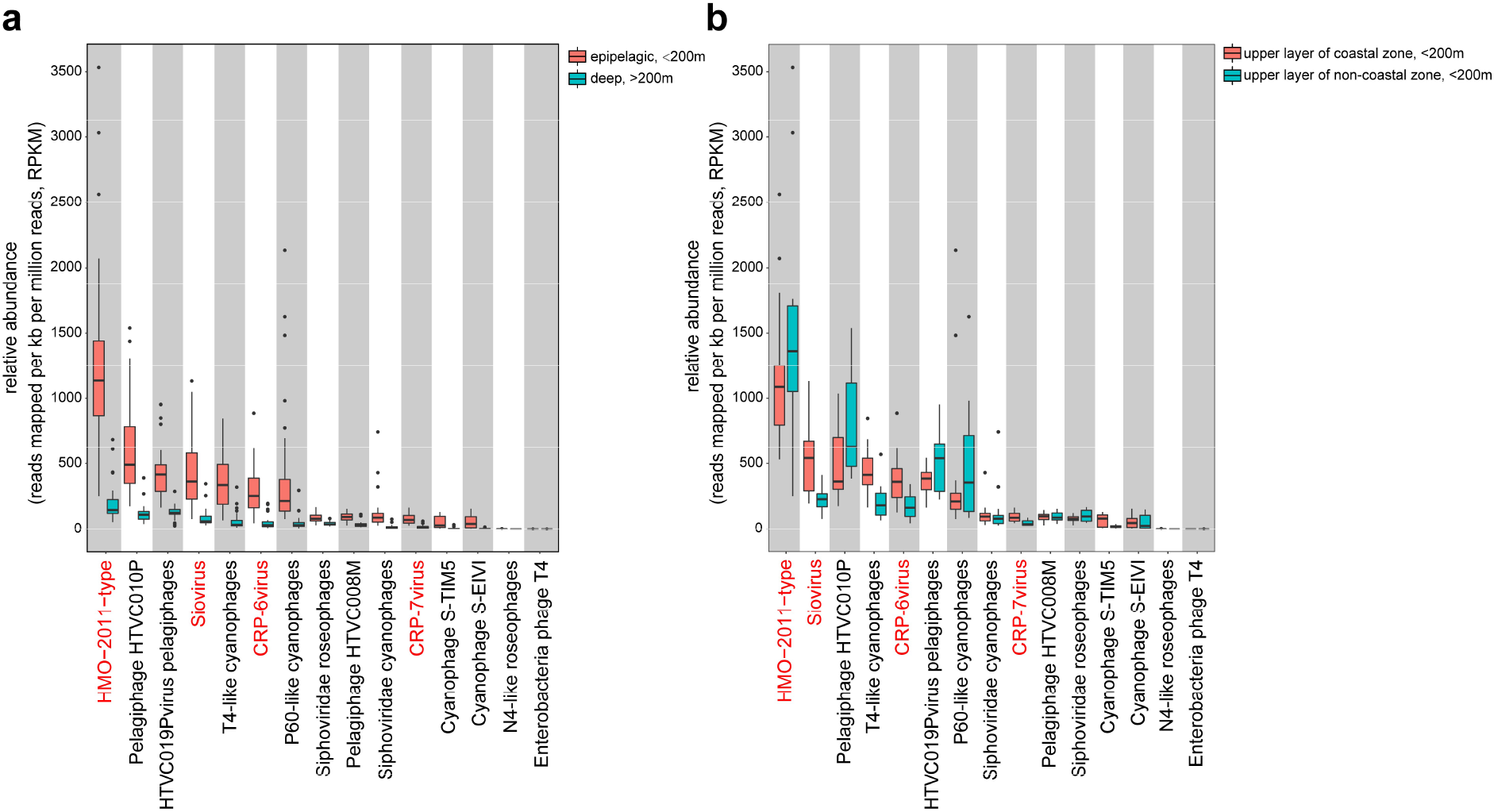
Box plots indicate the Rrelative abundance of major phage groups in marine viromes. Relative abundance (y axis) of each phage group (sorted according to its average relative abundance, x axis) in each virome dataset were calculated and normalized as RPKM (the number of reads recruited per kilobase pairs in average genome size per million of reads in the virome dataset). The marine virome datasets were from Pacific Ocean Virome (POV), India Ocean Virome (IOV), Scripps Pier virome (SPV) and Malaspina Expedition virome (ME). A Relative abundance of major phage groups in upper ocean samples (water depth <200m) compared with their abundance in deep ocean samples (water depth >200m). B. Relative abundance of major phage groups in coastal water samples compared with their abundance in non-coastal water samples. The phage groups containing RCA phages were shown in red.

In the *HMO-2011virus* genus, RCA phages CRP1 and CRP-2 cannot be well separated from HMO-2011 based on the genome content and sequence identity (Fig. 1b, 1c and Additional file 1: Figure S3). In most shared ORFs, CRP1 is more similar to HMO-2011 than to CRP-2 and CRP-3 (Additional file 1: Figure S3). The read recruitment analysis shows of the 97,684 reads assigned to the *HMO-2011virus* group in POV datasets, 78349, 74699 and 66560 reads could be mapped to the genome of HMO-2011, CRP-1 and CRP-2, respectively (Additional file 1: Figure S4). And the recruited reads mapped to the genomes of HMO-2011, CRP-1 and CRP-2 show highly similar distribution patterns of sequence identities and bitscores (Additional file 1: Figure S4 and Additional file 1: Figure S5). In addition, similar to a previous study [10], the majority of recruitments were phage genes associated with DNA metabolism and replication, structure and DNA packaging, which are conserved among CRP-1, CRP-2 and HMO-2011 (Additional file 1: Figure S4). Taken together, the results of our study suggest that it is difficult to separate the viromic reads assigned to the SAR116 phages and RCA phages within the *HMO-2011virus* genus.

We also performed the phylogenetic placement of the translated DNA polymerase sequences from POV datasets. In total, 5794 POV reads assigned to *HMO-2011virus* DNA polymerase and CRP-3 DNA polymerase were placed to the DNA polymerase reference tree (Fig. 5). The DNA polymerase gene phylogeny reveals that HMO-2011-type group contains remarkably diverse subgroups (Fig. 5), and a significant fraction of these DNA polymerase reads were placed near the reference sequence for HMO-2011, CRP-1 and CRP-2 (Fig. 5), suggesting that SAR116 and RCA roseobacters are probably two most important hosts for HMO-2011-type phages.

**Fig 5.**
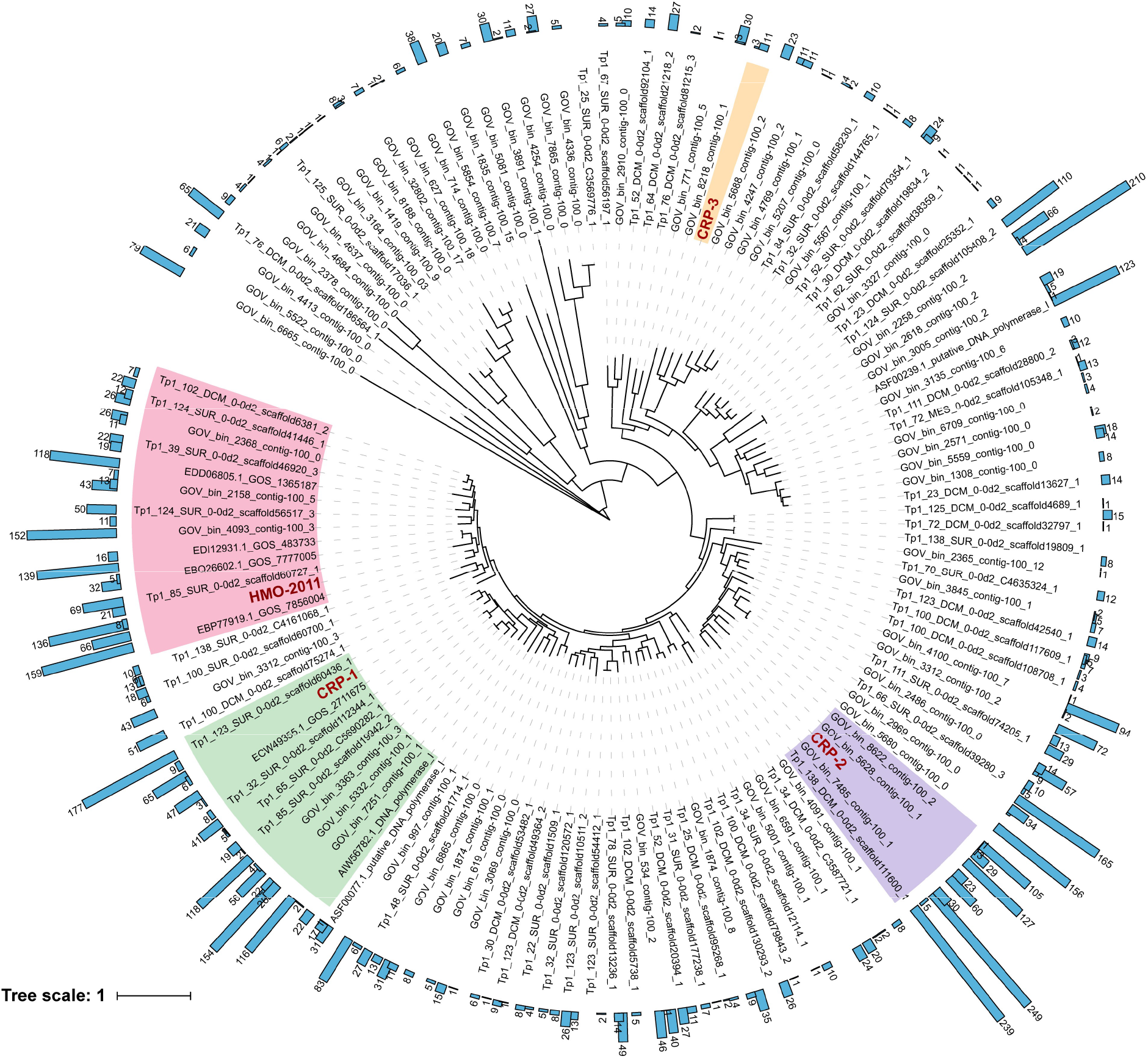
Phylogenetic placement of POV reads with homology to the DNA polymerase of HMO-2011virus phage. 5794 metagenomic reads were placed within the reference tree containing diverse homologs of HMO-2011-type DNA polymerase. HMO-2011 subgroup, CRP-1 subgroup, CRP-2 subgroup and are CRP-3 subgroup indicated in pink, cyan, purple and yellow, respectively (≥70% aa identity within group). The numbers of mapped viromic reads were shown by blue bars.

CRP-4 and CRP-5 are affiliated with the *Siovirus* genus which is widely distributed and is among the top most abundant phage groups in marine viromes. Roseophage SIO1 was previously found to be abundant in the POV datasets [9], and is ubiquitous in marine metagenomic datasets [38]. In our analyses, the results show that the relative abundance of *Siovirus* was comparable with those of pelagiphage *HTVC019Pvirus* and T4-like cyanophages in the upper ocean viomic datasets of POV, SPV and IOV (Fig.4a), and *Siovirus* was the second most abundant phage group in the coastal viromic datasets (Fig.4b). In GOV datasets, we observed that *Siovirus* exhibited significant lower abundance (Additional file 1: Figure S6).

The two new genera of RCA phages, *CRP-6virus* and *CRP-7virus*, are also ubiquitous in the ocean, as they were present in all marine viromic datasets. *CRP-6virus* was more abundant than *CRP-7virus* in almost all cases (Fig. 4, Additional file 1: Figure S6). In the upper layer of coastal zone, the relative abundance of *CRP-6virus* phages were comparable with *HTVC019Pvirus* pelagiphages and T4-like cyanophages (Fig. 4). Among these four RCA phage groups, *CRP-7virus* was the least present in the marine viromes but was still abundant and widely distributed in the ocean. The relative abundance of *CRP-7virus* was higher than those of N4-like roseophages and roseosipviruses in the vast majority of marine viromes (Fig. 4, Additional file 1: Figure S6).

Interestingly, except for HMO-2011-type, other phage groups presented by RCA phages exhibited significant higher abundance in the coastal waters than in the open ocean and intermediate regions (Fig.4, Additional file 1: Figure S6). In addition, all phage groups examined in this study were predominant in upper ocean viromes (<200 m) (Fig.4, Additional file 1: Figure S6). In congruence, roseobacters are reported predominate in the coast of temperate and polar environments [17, 18, 20–23], suggesting the tight co-occurrence and co-corelevant dynamic of RCA and RCA phages.

Our analysis clearly reveals the significant contribution of RCA phages to the diversity and abundance of marine viruses. As inferred from the “Kill the Winner” hypothesis, members of abundant RCA *Roseobacter* are subjected to a high level of density-dependent phage predation pressure. Therefore, it is not surprising that RCA phages constitute an important component of marine viral communities.

### Phage-host strategies

Oceans contain highly abundant viruses, many of which are still waiting to be discovered. Similar to SAR11, RCA represents a type of bacteria with reduced genome and oligotrophic adaptation [26]. It is intriguing that slow-growing heterotrophic bacteria such as SAR11 [9, 52], SAR116 [10] and RCA, are susceptible to infection by diverse podoviruses. Furthermore, these podoviruses are always highly ranked with respect to viromic fragment recruitment analyses (Fig. 4). Although many podoviruses have also been isolated from fast-growing roseobacters, such as *Ruegeria pomerioyi* DSS-3, *Sulfitobacter* spp. and *Dinoroseobacter shibae* DFL 12 [27, 28, 30, 31], most of the existing roseopodoviruses are N4-like phages, which were the least present in marine viromes (Fig. 4) despite that they infect multiple fast-growing *Roseobacter* lineages. Podoviruses are known to be lytic and have narrow host range in general. The high abundance and diversity of podoviruses that infect SAR11, SAR116 and RCA bacteria suggests a unique phage-host strategy between the podoviruses and K-selected bacteria (slow-growing but abundant) [11, 53, 54]. This type of “Kill-the-Winners” strategy could be important to maintain the equilibrium of K-selected or specialist bacteria in the oceans.

Many easy-to-grow roseobacters are generalists (utilize a variety of organic compounds) and have relatively larger genomes than RCA species [26]. These bacteria can quickly respond to environmental changes and are considered to be r-selected bacteria (fast-growing but sporadically abundants) [53, 54]. The acquisition of new genetic features is crucial for niche adaptation of r-selected roseobacters. Prophages are common in r-selected bacteria, while few or no prophages were found in K-selected bacteria [14, 26, 53, 55]. Many siphoviruses have been isolated from the fast-growing roseobacters [27], and it is possible that these siphoviruses play a key role in horizontal gene transfer in fast-growing roseobacters. On the other hand, K-selected bacteria display reduced genomes and oligotrophic adaptation, utilizing a narrow range of organic compounds (specialists). Prophage are rarely found in K-selected bacteria. Although some podoviruses infecting SAR11, SAR116 and RCA bacteria contain an integrase [9, 10, 52], the relatively low content of integration sequences of *HTVC019Pvirus* pelagiphages in Global Ocean Sampling (GOS) metagenomic database suggested that these phages infrequently lysogenize their hosts [52].

## Conclusions

Here, we first reported and described seven phages that infect an important group of marine bacteria-RCA *Roseobacter* lineage. RCA represents a group of difficult-to-culture, relatively slow-growing, but abundant bacterioplankton. The successful isolation and cultivation of the RCA strains in the laboratory facilitated the discovery of novel roseophage groups. The seven RCA phage genomes elucidated in this study contribute to the expansion of the diversity of cultured marine roseophages. It is surprising to identify such diverse groups of podoviruses that infect these closely related RCA strains. Marine viral communities contain numerous novel phage types without representative isolates. Given the ecological significance of their hosts and their ubiquity in the ocean, our study provides valuable experimental model systems for investigating phage ecology and phage-host interactions.

The three RCA strains used in this study grow slowly in diluted medium and do not reach high cell densities compared to fast-growing roseobacters. Because many groups of abundant bacteria and archaea have not been cultivated, their interactions with viruses are still unknown. We advocate that additional efforts are needed to isolate phages that infect abundant but not-yet-cultivated bacterioplankton to better understand the viral diversity in the ocean and interpret marine viromic datasets.

## Methods

### Cultivation of RCA strains, bacterial DNA extraction and PCR

The *Roseobacter* strains FZCC0023, FZCC0040, FZCC0042 and FZCC0083 were isolated on the 13^th^ of May, 2017 from the coastal water of Pingtang island in China (lat.’ N25°26’, long. E119°47’) using the dilution-to-extinction method [16]. All strains were grown in an autoclaved seawater based medium with excess vitamins [56] and amended with 1 mM NH4Cl, 100 μM KH_2_PO_4_, 1 μM FeCl_3_, and mixed carbon source [57]. Cultures were incubated at 20 °C. Genomic DNA of the RCA strains was extracted using a DNeasy Blood & Tissue Kit (Qiagen, Valencia, CA, USA) following the manufacturer’s protocol. The16S rRNA genes were amplified by PCR using the primers 16S-27F and 16S-1492R [58]. The primers 16S-907F and 23S-189R were used for PCR amplification of 16S-23S rDNA intergenic transcribed spacer (ITS) sequences [59, 60]. The 16S rRNA gene and ITS sequences of three strains were obtained by Sanger sequencing and assembled using ChromasPro (Technelysium Pty. Ltd., Tewantin QLD, Australia). The 16S rRNA and ITS sequences have been deposited in the GenBank database under the accession numbers MK335922 to MK335924.

### Source waters and RCA phage isolation

The water samples used to isolate RCA phages were collected from a variety of coastal stations (Table 1). The seawater samples were filtered through 0.1 μm pore-size filters and stored at 4 °C prior to use. The phages were isolated according to procedure detailed in a previous report [9]. Briefly, 0.1μm filtered water samples were inoculated with exponentially-growing host cultures and monitored for cell lysis using a Guava EasyCyte cell counter (Merck Millipore, Billerica, MA, USA). For cell lysis cultures, the presence of phage particles was further confirmed by epifluorescence microscopy with SYBR Green I (Invitrogen, Eugene, OR, USA) [61]. Purified RCA phage clones were obtained using the dilution-to-extinction method. The purity of the RCA phages was assessed by genome sequencing.

### Transmission electron microscopy

RCA phage lysates were filtered through 0.1μm pore-size filters and then concentrated using Amicon Ultra Centrifugal Filters (30-kDa, Merck Millipore). Concentrated phages were absorbed onto copper grids in the dark and stained with 2% uranyl acetate for two minutes. Samples were observed using a Hitachi transmission electron microscope at 80 kV.

### Cross-infection experiments

The cross-infectivity of seven RCA phages was tested in liquid medium against three RCA strains in duplicate. Exponentially-growing cultures of three RCA strains were incubated with each RCA phage at a phage-to-host ratio of approximately 20. Cell lysis was monitored by using a Guava EasyCyte Flow Cytometer and phage particles were enumerated by epifluorescence microscopy.

### Phage DNA preparation, genome sequencing

250 ml phage lysates were filtered through 0.1 μm pore-size Supor membrane and further concentrated using 30 kDa MW cutoff Amicon Ultra Centrifugal Filters. Phage DNA was extracted using the formamide extraction method [62]. The genomes of the RCA phages were sequenced using Illumina paired-end HiSeq 2500 sequencing approach at Beijing Mega Genomics Technology (Beijing, China). Quality filtration, removal of adapter and low quality sequences, *de novo* assembly of reads were performed using CLC Genomic Workbench 11.0.1 (QIAGEN, Hilden, Germany) with the default settings. The gaps in phage genomes were closed by PCR amplification of the regions between the contigs and followed by conventional Sanger sequencing.

### Genome annotation and Comparative Genomic analysis

Putative open reading frames (ORFs) longer than 120 bp were predicted from assembled RCA phage genomes by GeneMark [63], RAST server [64], and manual inspection. tRNAs were detected using tRNAscan-SE [65]. Putative biological functions were assigned to translated ORFs using BLASTp (amino acid identity ≥30%, alignment coverage ≥50%, and E-value ≤1E-3) against the NCBI non-redundant (nr) database and the NCBI-RefSeq database in comparison with known proteins. PFAM and HHpred were also used to identify the protein families and distant protein homologs, respectively. The genome sequences of RCA phages have been deposited in the GenBank database under the accession numbers MK613343 to MK613349.

### Determination of CRP-3 integration sites

The integration sites of CRP-3 were identified by following a strategy described in a previous study [52]. Briefly, DNA of phage-infected cells was sequenced using the Illumina HiSeq 2500. The raw reads were quality filtered, trimmed and mapped to the CRP-3 genome using CLC Genomic Workbench 11.0.1. Sequences that mapped to the CRP-3 genome were manually inspected to detect the phage-host hybrid sequences. The resulting hybrid sequences were analysed to identify the integration sites and their locations on the host genomes. PCR primer sets (attL-F: TTCGGGACTGGAAGCATAC, attL-R: CCTAAAGGCAGGAGGATACAC, attR-F: CAGAGCCTCTTTGTGATGGTC and attR-R: AGGCACACTGGACTATACACAG) were designed based on the predicted *attL* and *attR* sites.

### Phylogenetic analysis

Phylogenetic trees based on 16S rRNA gene and internal transcribed spacer (ITS) sequences were constructed using FastTree 2.1 [66] with the following settings: FastTree -gtr -nt -boot 1000 Sequences < alignment_file > tree_file. Sequences were aligned using MAFFT [67]. Amino acid sequences of DNA polymerase were aligned and used for phylogenetic analysis. Alignments of DNA polymerase gene sequences were constructed with MUSCLE [68] and edited with Gblocks [69]. Alignments were evaluated for the optimal amino acid substitution models using ProtTest [70], and maximum likelihood tree was constructed by RAxML v8 [71].

### Network Analysis

Ninety-eight complete *Podoviridae* genomes were downloaded from the National Center for Biotechnology Information (NCBI) (Additional files2: Table S1). All proteins were compared using all-versus-all BLASTp (e-value ≤1E-5, bitscore ≥50, amino acid identity ≥30%), after which protein clusters were defined using the Markov clustering algorithm (MCL) [72]. The similarity between two phages based on the number of shared protein families was calculated using the hypergeometric formula that has been described in detail in a previous reports: 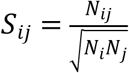, where *N_ij_* is the numbers of shared protein families between phage i and phage j, and *N_i_* and *N_j_* are the numbers of proteins in phage i and phage j, respectively [73]. The network was visualized using Cytoscape. In the network, nodes represent phages and the weight of each edges represent the distance between two phages based on the similarity, with a cutoff of 0.05.

### Phylogenetic placement of viromic reads assigned to HMO-2011-type DNA polymerase genes

Amino acid sequences of HMO-2011-type DNA polymerase homologs were extracted from viral single-amplified genomes (vSAGs), Global Ocean Sampling (GOS) metagenomic sequences and assembled Global Oceanic Virome (GOV) viral populations using BLASTp (e-value ≤10^-3^, coverage ≥80%). 129 nearly full-length environmental viral DNA polymerase sequences were aligned with HMO-2011-type DNA polymerase sequences using MAFFT [67]. Phylogenetic tree of this alignment was constructed by RAxML v8 [71] with the following settings: raxmlHPC-PTHREADS -T 12 -f a -x 12345 -m PROTGAMMAWAGF -s alignment_file -# 100 - p 12345. The Pacific Ocean Virome (POV) reads assigned to the DNA polymerase genes of the *HMO-2011virus* phages were translated and placed on the phylogenetic tree using the pplacer v1.1 [74]. The resulting tree phylogeny was visualized and manipulated using iTOL v4 [75].

### RCA phages fragment recruitment from marine virome

Virome datasets from the Pacific Ocean Virome (POV), Scripps Pier virome (SPV), India Ocean Virome (IOV) and Malaspina Expedition virome (ME) were used for phage reciprocal metagenomic fragment recruitment analysis (Additional files2: Table S3). The analysis was performed according to procedure detailed in a previous report [9], the detailed steps are as follows:

Each of the virome reads was searched as a query again the RefSeq viral database (release 88), 11 newly sequenced *HTVC019Pvirus* pelagiphages [52], and seven RCA phage genomes in this study using DIAMOND BLASTx with an e-value cutoff of 10^-3^ and a bitscore cutoff 40.

1. The resulting hits were extracted from the virome datasets and used as queries for BLASTx, against a protein database containing the following:

a. A subset of the viral genomes from RefSeq, excluding 2b.
b. The viral genomes used in this study (Additional files2: Table S4), including the 7 newly sequenced RCA phage genomes.
c. A subset of the bacterial genomes from RefSeq (release 81).
2. Metagenomic sequences that returned a best-hit of the query genome from (2b) were identified and extracted from the metagenomic datasets.
3. The relative abundances of each phage group were calculated and normalized as the number of reads recruited to the group divided by the total number of kilobase pairs in average genome size and divided by the total number of million reads the virome dataset (reads mapped per kb of genome sequence per million of virome reads, RPKM).

Due to the large amount of sequencing data in the Global Oceans Viromes (GOV) datasets (>500G), a different strategy was used to determine the relative abundances of different phage groups in GOV. GOV reads were recruited onto the phage genomes (Additional files2: Table S4) using BLASTx with an e-value cutoff of 10^-10^. If a read was recruited to more than one phage genome, the read was associated with the phage that provided the highest bitscore.

## Declarations

### Ethics approval and consent to participate

Not applicable

### Consent for publication

Not applicable

### Availability of data and materials

The 16S rRNA and ITS sequences have been deposited in the GenBank database under the accession numbers MK335922 to MK335924. The genome sequences of RCA phages have been deposited in the GenBank database under the accession numbers MK613343 to MK613349.

### Author contributions

ZFZ, FC and YZ conceived and designed the project and the experiments. FQ. and MY isolated the phages. FQ and JS prepared the samples for transmission electron microscopy. ZFZ and ZZ performed the cross-infection experiment. YZ and ZFZ performed the bioinformatic analyses. XC and HZ performed the metagenomic fragment recruitment analyses. FC and HL gave technical support and conceptual advice. ZFZ, FC and Y.Z wrote the paper.

## Acknowledgements

We thank Jie Wang for providing the water samples. We also thank Chen Li for her assistance in TEM.

## Additional files

**Additional files1: Figure S1** Phylogenetic relationship between three RCA isolates and the representatives belonging to the *Roseobacter* clade of the *Alphaproteobacteria*. **Figure S2** Integration of CRP-3 in the FZCC0040 genome. **Figure S3** Percent amino acid identity of phage HMO-2011, CRP-2 and CRP-3 to the CRP-1 ORFs. **Figure S4** Fragment recruitment plot of % amino acid identity and bitscores of Pacific Ocean Virome reads against the ORFs of phage HMO-2011, CRP-1 and CRP-2. **Figure S5** Boxplot distributions of % amino acid identity and bitscores of the POV reads assigned to *HMO-2011virus* group against each *HMO-2011virus* genome. **Figure S6** Relative abundance of major phage groups in Global Oceans Viromes (GOV).

**Additional files2: Table S1** List of phage genomes included in network analysis. **Table S2** Summary of putative AMG genes and genes involved in other cellular processes content across seven RCA phages. **Table S3** The list of marine virome datasets analyzed in this study. **Table S4** List of all marine phage genomes included in the metagenome analysis.

**Additional files3: Dataset 1** Predicted ORFs of seven RCA phages

## References

1. Wommack KE, Colwell RR. Virioplankton: viruses in aquatic ecosystems. Microbiol Mol Biol Rev. 2000;64:69–114.

2. Suttle CA. Marine viruses--major players in the global ecosystem. Nat Rev Microbiol. 2007;5:801–12.

3. Fuhrman JA. Marine viruses and their biogeochemical and ecological effects. Nature. 1999;399:541–48.

4. Kogure K, Simidu U, Taga N. A tentative direct microscopic method for counting living marine bacteria. Can J Microbiol. 1979;25:415–20.

5. Mizuno CM, Rodriguezvalera F, Kimes NE, Ghai R. Expanding the marine virosphere using metagenomics. PLoS Genet. 2013;9:e1003987.

6. Roux S, Hallam SJ, Woyke T, Sullivan MB. Viral dark matter and virus–host interactions resolved from publicly available microbial genomes. eLife. 2015;4:e08490.

7. Brum JR, Ignacioespinoza JC, Roux S, Doulcier G, Acinas SG, Alberti A, et al. Patterns and ecological drivers of ocean viral communities. Science. 2015;348:1261498.

8. Roux S, Brum JR, Dutilh BE, Sunagawa S, Duhaime MB, Loy A, et al. Ecogenomics and potential biogeochemical impacts of globally abundant ocean viruses. Nature. 2016;537:689–93.

9. Zhao Y, Temperton B, Thrash JC, Schwalbach MS, Vergin KL, Landry ZC, et al. Abundant SAR11 viruses in the ocean. Nature. 2013;494:357–60.

10. Kang I, Oh HM, Kang D, Cho JC. Genome of a SAR116 bacteriophage shows the prevalence of this phage type in the oceans. Proc Natl Acad Sci USA. 2013;110:12343–48.

11. Giovannoni SJ. SAR11 Bacteria: The most abundant plankton in the oceans. Ann Rev Mar Sci. 2017;9:231–55.

12. Rappé MS, Kemp PF, Giovannoni SJ. Phylogenetic diversity of marine coastal picoplankton 16S rRNA genes cloned from the continental shelf off Cape Hatteras, North Carolina. Limnol Oceanogr. 1997;42:811–26.

13. Giovannoni SJ, Vergin KL. Seasonality in ocean microbial communities. Science. 2012;335:671–6.

14. Grote J, Bayindirli C, Bergauer K, Carpintero de Moraes P, Chen H, D’Ambrosio L, et al. Draft genome sequence of strain HIMB100, a cultured representative of the SAR116 clade of marine *Alphaproteobacteria*. Stand Genomic Sci. 2011;5:269–78.

15. Rappe MS, Connon SA, Vergin KL, Giovannoni SJ. Cultivation of the ubiquitous SAR11 marine bacterioplankton clade. Nature. 2002;418:630–3.

16. Stingl U, Tripp HJ, Giovannoni SJ. Improvements of high-throughput culturing yielded novel SAR11 strains and other abundant marine bacteria from the Oregon coast and the Bermuda Atlantic Time Series study site. ISME J. 2007;1:361–71.

17. Wagner-Dobler I, Biebl H. Environmental biology of the marine *Roseobacter* lineage. Annu Rev Microbiol. 2006;60:255–80.

18. Brinkhoff T, Giebel HA, Simon M. Diversity, ecology, and genomics of the *Roseobacter* clade: a short overview. Arch Microbiol. 2008;189:531–39.

19. Luo H, Loytynoja A, Moran MA. Genome content of uncultivated marine *Roseobacters* in the surface ocean. Environ Microbiol. 2012;14:41–51.

20. Zhang Y, Sun Y, Jiao N, Stepanauskas R, Luo H. Ecological genomics of the uncultivated marine *Roseobacter* lineage CHAB-I-5. Appl Environ Microbiol. 2016;82:2100–11.

21. Giebel HA, Kalhoefer D, Lemke A, Thole S, Gahl-Janssen R, Simon M, et al. Distribution of *Roseobacter* RCA and SAR11 lineages in the North Sea and characteristics of an abundant RCA isolate. ISME J. 2011;5:8–19.

22. Selje N, Simon M, Brinkhoff T. A newly discovered *Roseobacter* cluster in temperate and polar oceans. Nature. 2004;427:445–8.

23. Giebel HA, Brinkhoff T, Zwisler W, Selje N, Simon M. Distribution of *Roseobacter* RCA and SAR11 lineages and distinct bacterial communities from the subtropics to the Southern Ocean. Environ Microbiol. 2009;11:2164–78.

24. Mayali X, Franks PJ, Azam F. Cultivation and ecosystem role of a marine *Roseobacter* clade-affiliated cluster bacterium. Appl Environ Microbiol. 2008;74:2595–603.

25. Giebel HA, Kalhoefer D, Gahl-Janssen R, Choo YJ, Lee K, Cho JC, et al. *Planktomarina temperata* gen. nov., sp. nov., belonging to the globally distributed RCA cluster of the marine *Roseobacter* clade, isolated from the German Wadden Sea. Int J Syst Evol Microbiol. 2013;63:4207–17.

26. Voget S, Wemheuer B, Brinkhoff T, Vollmers J, Dietrich S, Giebel HA, et al. Adaptation of an abundant *Roseobacter* RCA organism to pelagic systems revealed by genomic and transcriptomic analyses. ISME J. 2015;9:371–84.

27. Zhan Y, Chen F. Bacteriophages that infect marine roseobacters: genomics and ecology. Environ Microbiol. 2018; in press.

28. Rohwer F, Segall A, Steward G, Seguritan V, Breitbart M, Wolven F, et al. The complete genomic sequence of the marine phage Roseophage SIO1 shares homology with nonmarine phages. Limnol Oceanogr. 2000;45:408–18.

29. Zhao Y, Wang K, Jiao N, Chen F. Genome sequences of two novel phages infecting marine roseobacters. Environ Microbiol. 2009;11:2055–64.

30. Ankrah NY, Budinoff CR, Wilson WH, Wilhelm SW, Buchan A. Genome sequences of two temperate phages, ΦCB2047-A and ΦCB2047-C, infecting *Sulfitobacter* sp. strain 2047. Genome Announc. 2014;2:e00108–14.

31. Li B, Zhang S, Long L, Huang S. Characterization and complete genome sequences of three N4-Like roseobacter phages isolated from the South China Sea. Curr Microbiol. 2016;73:409–18.

32. Yang Y, Cai L, Ma R, Xu Y, Tong Y, Huang Y, Jiao N, Zhang R, et al. A novel roseosiphophage isolated from the oligotrophic South China Sea. Viruses. 2017;9:109.

33. Lavigne R, Seto D, Mahadevan P, Ackermann HW, Kropinski AM. Unifying classical and molecular taxonomic classification: analysis of the Podoviridae using BLASTP–based tools. Res Microbiol. 2008;159:406–14.

34. Labonte JM, Reid KE, Suttle CA. Phylogenetic analysis indicates evolutionary diversity and environmental segregation of marine podovirus DNA polymerase gene sequences. Appl Environ Microbiol. 2009;75:3634–40.

35. Meier-Kolthoff JP, Goker M. VICTOR: genome-based phylogeny and classification of prokaryotic viruses. Bioinformatics. 2017;33:3396–404.

36. Chafee M, Fernandez-Guerra A, Buttigieg PL, Gerdts G, Eren AM, Teeling H, et al. Recurrent patterns of microdiversity in a temperate coastal marine environment. ISME J. 2018;12:237–52.

37. Kang I, Jang H, Oh HM, Cho JC. Complete genome sequence of *Celeribacter* bacteriophage P12053L. J Virol. 2012;86:8339–40.

38. Bischoff V, Bunk B, Meier-Kolthoff JP, Sproer C, Poehlein A, Dogs M, et al. Cobaviruses - a new globally distributed phage group infecting *Rhodobacteraceae* in marine ecosystems. ISME J. 2019; in press.

39. Huang S, Zhang S, Jiao N, Chen F. Comparative genomic and phylogenomic analyses reveal a conserved core genome shared by estuarine and oceanic cyanopodoviruses. PLoS One. 2015;10:e0142962.

40. Holmfeldt K, Solonenko N, Shah M, Corrier K, Riemann L, Verberkmoes NC, et al. Twelve previously unknown phage genera are ubiquitous in global oceans. Proc Natl Acad Sci USA. 2013;110:12798–803.

41. Breitbart M, Thompson LR, Suttle CA, Sullivan MB. Exploring the vast diversity of marine viruses. Oceanogr. 2007;20:135–9.

42. Murphy J, Mahony J, Ainsworth S, Nauta A, Van Sinderen D. Bacteriophage orphan DNA methyltransferases: insights from their bacterial origin, function, and occurrence. Appl Environ Microbiol. 2013;79:7547–55.

43. Bryan MJ, Burroughs NJ, Spence EM, Clokie MR, Mann NH, Bryan SJ. Evidence for the intense exchange of MazG in marine cyanophages by horizontal gene transfer. PLoS ONE. 2008;3:e2048.

44. Angly F, Youle M, Nosrat B, Srinagesh S, Rodriguez-Brito B, McNairnie P, et al. Genomic analysis of multiple Roseophage SIO1 strains. Environ Microbiol. 2009;11:2863–73.

45. Duhaime MB, Wichels A, Waldmann J, Teeling H, Glöckner FO. Ecogenomics and genome landscapes of marine *Pseudoalteromonas* phage H105/1. ISME J. 2011;5:107–21.

46. Lee S, Kim MH, Kang BS, Kim JS, Kim GH, Kim YG, et al. Crystal structure of *Escherichia coli* MazG, the regulator of nutritional stress response. J Biol Chem. 2008;283: 15232–40.

47. Gross M, Marianovsky I, Glaser G. MazG-a regulator of programmed cell death in Escherichia coli. Mol Microbiol. 2010;59:590–601.

48. Grant CM. Role of the glutathione/glutaredoxin and thioredoxin systems in yeast growth and response to stress conditions. Mol Microbiol. 2001;39:533–41.

49. Lillig CH, Berndt C, Holmgren A. Glutaredoxin systems. Biochim Biophys Acta 2008;1780:1304–17.

50. Salah Ud-Din AI, Tikhomirova A, Roujeinikova A. Structure and functional diversity of GCN5-Related N-Acetyltransferases (GNAT). Int J Mol Sci. 2016;17:1018.

51. Pattridge KA, Weber CH, Friesen JA, Sanker S, Kent C, Ludwig ML. Glycerol-3-phosphate cytidylyltransferase. J. Biol. Chem. 2003;278:51863.

52. Zhao Y, Qin F, Zhang R, Giovannoni SJ, Zhang Z, Sun J, et al. Pelagiphages in the *Podoviridae* family integrate into host genomes. Environ Microbiol. 2018; e-pub ahead of print 26 November 2018; in press.

53. Lauro FM, McDougald D, Thomas T, Williams TJ, Egan S, Rice S, DeMaere MZ, et al. The genomic basis of trophic strategy in marine bacteria. Proc Natl Acad Sci USA. 2009;106:15527–33.

54. Yooseph S, Nealson KH, Rusch DB, McCrow JP, Dupont CL, Kim M, et al. Genomic and functional adaptation in surface ocean planktonic prokaryotes. Nature. 2010;468:60–6.

55. Oh HM, Kwon KK, Kang I, Kang SG, Lee JH, Kim SJ, et al. Complete genome sequence of “*Candidatus Puniceispirillum marinum*” IMCC1322, a representative of the SAR116 clade in the *Alphaproteobacteria*. J Bacteriol. 2010;192:3240–41.

56. Sun J, Steindler L, Thrash JC, Halsey KH, Smith DP, Carter AE, et al. One carbon metabolism in SAR11 pelagic marine bacteria. PLoS ONE. 2011;6:e23973.

57. Cho JC, Giovannoni SJ. Cultivation and growth characteristics of a diverse group of oligotrophic marine *Gammaproteobacteria*. Appl Environ Microbiol. 2004;70:432–40.

58. Lane DJ. 16S/23S rRNA sequencing. In Nucleic Acid Techniques in Bacterial. Edited by Systematics Stackebrandt E, Goodfellow M, John Wiley and Sons. New York: West Sussex; 1991:115–75.

59. Muyzer G, Smalla K. Application of denaturing gradient gel electrophoresis (DGGE) and temperature gradient gel electrophoresis (TGGE) in microbial ecology. Anton Leeuw. 1998;73:127–41.

60. Hunt DE, Klepac-Ceraj V, Acinas SG, Gautier C, Bertilsson S, Polz MF. Evaluation of 23S rRNA PCR primers for use in phylogenetic studies of bacterial diversity. Appl Environ Microbiol. 2006;72:2221–5.

61. Suttle CA, Fuhrman JA. Enumeration of virus particles in aquatic or sediment samples by epifluorescence microscopy. In Manual of Aquatic Viral Ecology. Edited by Wilhelm S, Weinbauer M, Suttle C,. American Society of Limnology and Oceanography: Waco, TX; 2010. 145–53.

62. Sambrook J, Russell DW. (Ed). Molecular Cloning: A Laboratory Manual. Cold Spring Harbor, New York: Cold Spring Harbor Laboratory; 2001.

63. Borodovsky M, McIninch J. GENMARK: Parallel gene recognition for both DNA strands. Comput Chem. 1993;17:123–33.

64. Aziz RK, Bartels D, Best AA, DeJongh M, Disz T, Edwards RA, et al. The RAST Server: rapid annotations using subsystems technology. BMC Genomics 2008;9:75.

65. Lowe TM, Eddy SR. tRNAscan-SE: a program for improved detection of transfer RNA genes in genomic sequence. Nucleic Acids Res. 1997;25:955–64.

66. Price MN, Dehal PS, Arkin AP. FastTree 2--approximately maximum-likelihood trees for large alignments. PLoS ONE. 2010;5:e9490.

67. Katoh K, Asimenos G, Toh H. Multiple alignment of DNA sequences with MAFFT. Methods Mol Biol. 2009;537:39–64.

68. Edgar RC. MUSCLE: multiple sequence alignment with high accuracy and high throughput. Nucleic Acids Res. 2004;32:1792–97.

69. Castresana J. Selection of conserved blocks from multiple alignments for their use in phylogenetic analysis. Mol Biol Evol. 2000;17:540–52.

70. Abascal F, Zardoya R, Posada D. ProtTest: selection of best-fit models of protein evolution. Bioinformatics. 2005;21:2104–05.

71. Stamatakis A. RAxML version 8: a tool for phylogenetic analysis and postanalysis of large phylogenies. Bioinformatics. 2014;30:1312–3.

72. Enright AJ, Van Dongen S, Ouzounis CA. An efficient algorithm for large-scale detection of protein families. Nucleic Acids Res. 2002;30:1575–84.

73. Iranzo J, Krupovic M, Koonin EV. The double-stranded DNA virosphere as a modular hierarchical network of gene sharing. mBio. 2016;7:e00978–16.

74. Matsen FA, Kodner RB, Armbrust EV. pplacer: linear time maximum-likelihood and Bayesian phylogenetic placement of sequences onto a fixed reference tree. BMC Bioinformatics. 2010;11:538.

75. Letunic I, Bork P. Interactive tree Of life (iTOL) v4: recent updates and new developments. Nucleic Acids Res. 2019; gkz239.

