## Supplementary Figures for "Diverse, abundant and novel viruses infecting “unculturable” but abundant marine bacteria"

**Additional files1:**

### Supplementary Figures for
### Diverse, abundant and novel viruses infecting "unculturable" but abundant marine bacteria

Zefeng Zhang[1], Feng Chen[2], Xiao Chu[3], Hao Zhang[3], Haiwei Luo[3], Fang Qin[1], Zhiqiang Zhai[1], Mingyu Yang[1], Jing Sun[4], Yanlin Zhao[1] *

[1] Fujian Provincial Key Laboratory of Agroecological Processing and Safety Monitoring, College of Life Sciences, Fujian Agriculture and Forestry University, Fuzhou, Fujian, 350002, China

[2] Institute of Marine and Environmental Technology, University of Maryland Center for Environmental Science, Baltimore, MD, USA

[3] Simon F. S. Li Marine Science Laboratory, School of Life Sciences and Partner State Key Laboratory of Agrobiotechnology, The Chinese University of Hong Kong, Shatin, Hong Kong SAR, China

[4] Yellow Sea Fisheries Research Institute, Chinese Academy of Fishery Sciences, Qingdao, Shandong, 266071, China

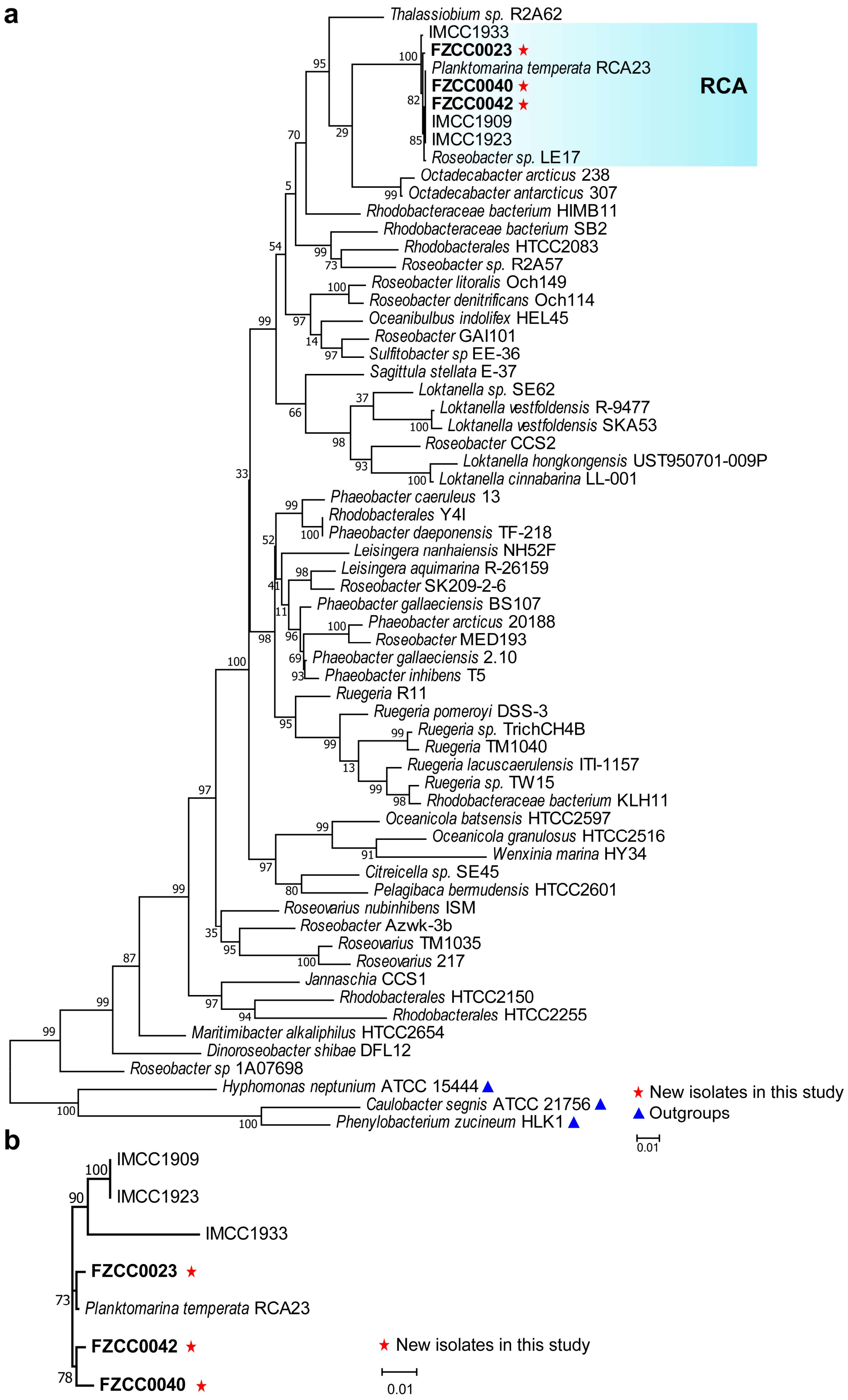


**Figure S1**: Phylogenetic relationship between three RCA isolates and the representatives belonging to the *Roseobacter* clade of the *Alphaproteobacteria*. **a** Phylogenetic analysis of 16S rRNA gene. **b** Phylogenetic analysis of our RCA isolates and four RCA representatives based on the ITS sequences.

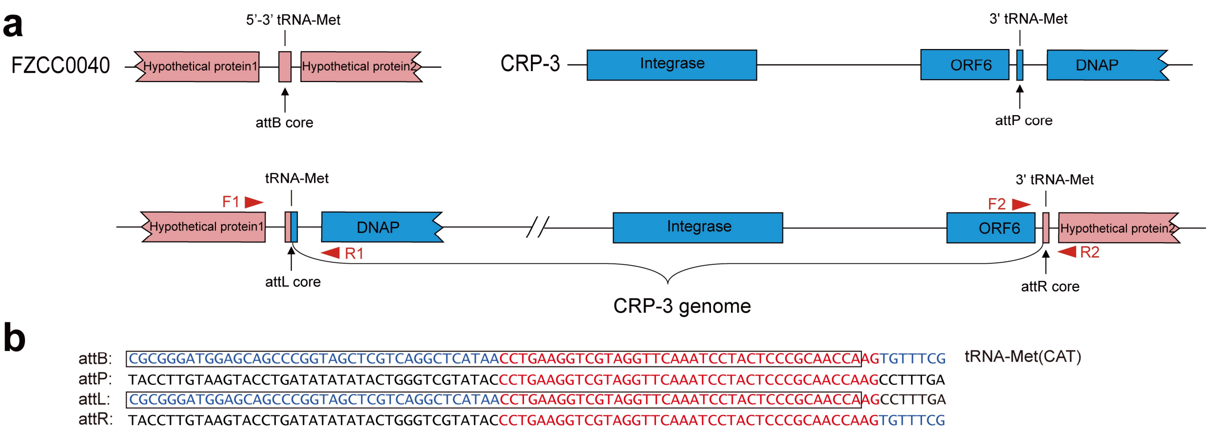


**Figure S2**: Integration of CRP-3 in the FZCC0040 genome. **a** CRP-3 and host FZCC0040 genes are shown in pink and blue respectively. The core sequences are indicated by the arrows. PCR primers are indicated by red triangles. **b** The DNA sequences of all integration sites. CRP-3 sequences, FZCC0040 sequences and the identical core sequences are shown in black, blue and red, respectively. The tRNA genes are boxed. Abbreviation: DNAP, DNA polymerase

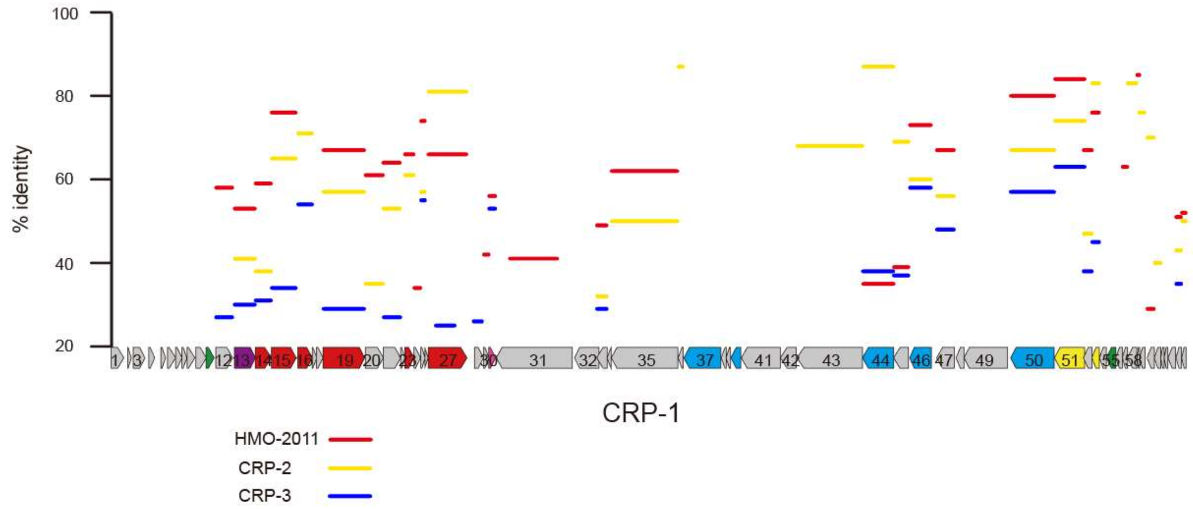


**Figure S3**: Percent amino acid identity of phage HMO-2011, CRP-2 and CRP-3 to the CRP-1 ORFs.

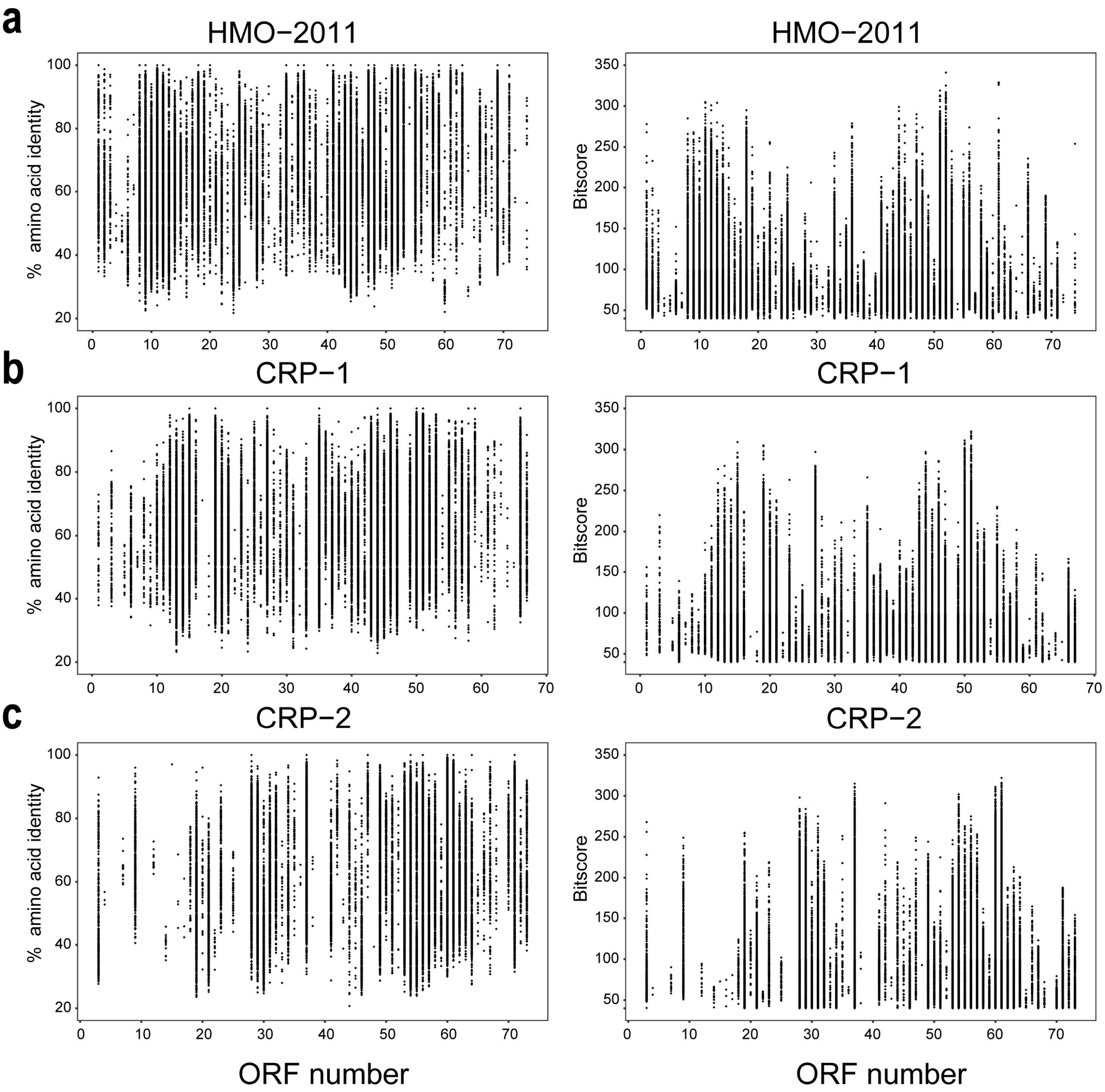


**Figure S4**: Fragment recruitment plot of % amino acid identity and bitscores of Pacific Ocean Virome reads against the ORFs of phage HMO-2011, CRP-1 and CRP-2. The total number of POV reads assigned to the *HMO-2011viruse* group is 97,684. Numbers of reads mapped to HMO-2011, CRP-1 and CRP-2 are 78,349, 74,699 and 66,560, respectively. **a,** HMO-2011. **b,** CRP-1. **c,** CRP-2

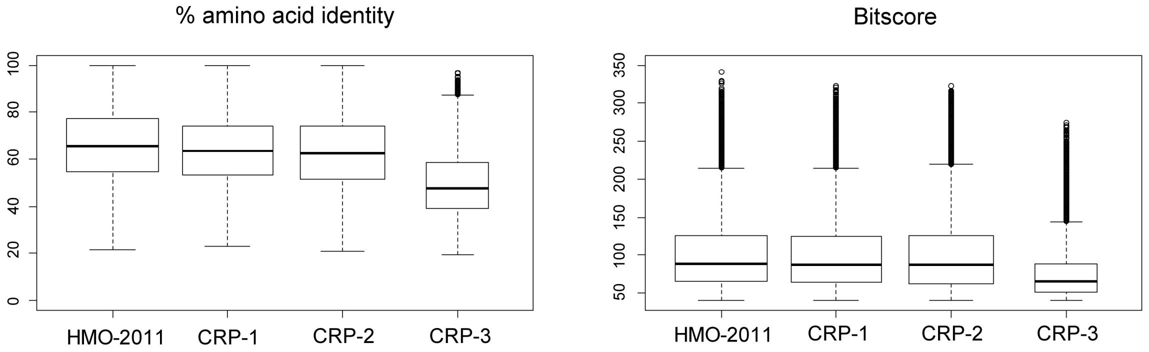


**Figure S5**: Boxplot distributions of % amino acid identity and bitscores of the POV reads assigned to *HMO-2011virus* group against each *HMO-2011virus* genome.

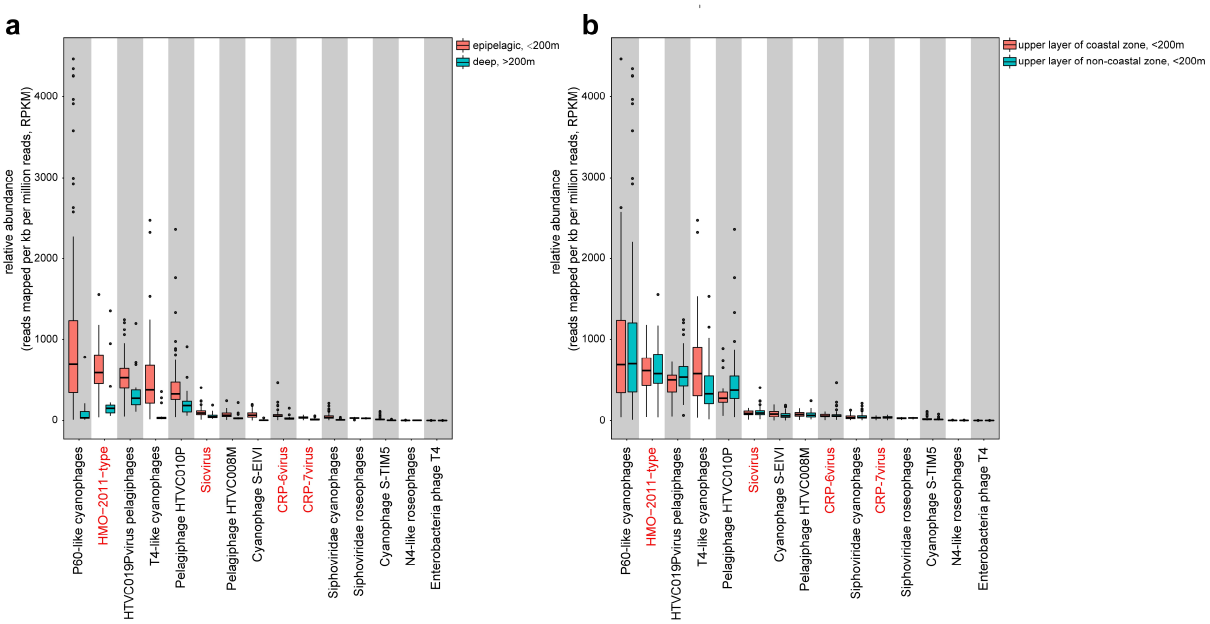


**Figure S6 Relative abundance of major phage groups in Global Oceans Viromes (GOV).** Relative abundance (y axis) of each phage group (sorted according to its median relative abundance, x axis) in each virome dataset were calculated and normalized as RPKM (the number of reads recruited per kilobase pairs in average genome size per million of reads in the virome dataset). A Relative abundance of major phage groups in upper ocean samples (water depth <200m) compared with their abundance in deep ocean samples (water depth >200m). B. Relative abundance of major phage groups in coastal water samples compared with their abundance in non-coastal water samples. The phage groups containing RCA phages were shown in red.
